# MICRORNAS REGULATED BY PREGNANCY TARGET THE HIV INTERACTOME

**DOI:** 10.1101/2025.07.21.665982

**Authors:** P.F.T. Cezar-de-Mello, J. Dreyfuss, P. Chen, H. Yamamoto, X. Gao, H. Pan, C. Morrison, G. F. Doncel, R. Barbieri, R. Fichorova

## Abstract

Innate immunity predictors of HIV risk are influenced by reproductive hormone levels, pregnancy, and lactation status. However, the molecular mechanisms underlying these associations remain unclear. MicroRNAs (miRNAs), as post-transcriptional regulators, play key roles in immune regulation and host-virus interactions. We hypothesized that physiological adaptations to pregnancy include the modulation of systemic miRNA expression, which in turn regulates host genes involved in HIV interaction, potentially influencing susceptibility during pregnancy. To test this, we leveraged a large longitudinal cohort from Uganda and Zimbabwe and analyzed 174 serum samples from 88 participants in pre-pregnancy (PP), pregnancy (P), and breastfeeding (BF) states using the HTG EdgeSeq platform. Differentially expressed (DE) miRNAs in pregnancy were identified with false discovery rate (FDR) < 0.1 (29 upregulated and 131 downregulated) by intersecting pairwise comparisons (P vs. PP and P vs. BF). Validated gene targets (2,733) were identified via miRWalk and modular enrichment analysis performed using Cytoscape/ClueGO. Enriched pathways (FDR<0.05) included Adaptive Immune Response, Hippo Signaling, Cellular Senescence, HSV-1 Infection, and two cancer-related pathways. Genes from pregnancy-enriched pathways overlapped with the known HIV-host interactome at the range of 37 to 88%. From this overlap, the Maximal Clique Centrally (MCC) score (cytoHubba) identified 47 unique hub genes acting as essential regulatory nodes of protein-protein interaction (PPI) network. These hub genes were further examined by their expression patterns predicted by DE miRNAs, and explored for their interactions with the HIV interactome, revealing connectivity with 18 HIV proteins - highest with Tat and gp120 - and impact on the HIV replication process. HLA-A was the most connected host gene, followed by BCL2L1, EIF2AK2, PTEN, and CYCS. These findings support the hypothesis that pregnancy-driven systemic miRNAs may shape HIV susceptibility by regulating hub genes with central role in viral evasion of host immunity.

**GRAPHIC ABSTRACT:** 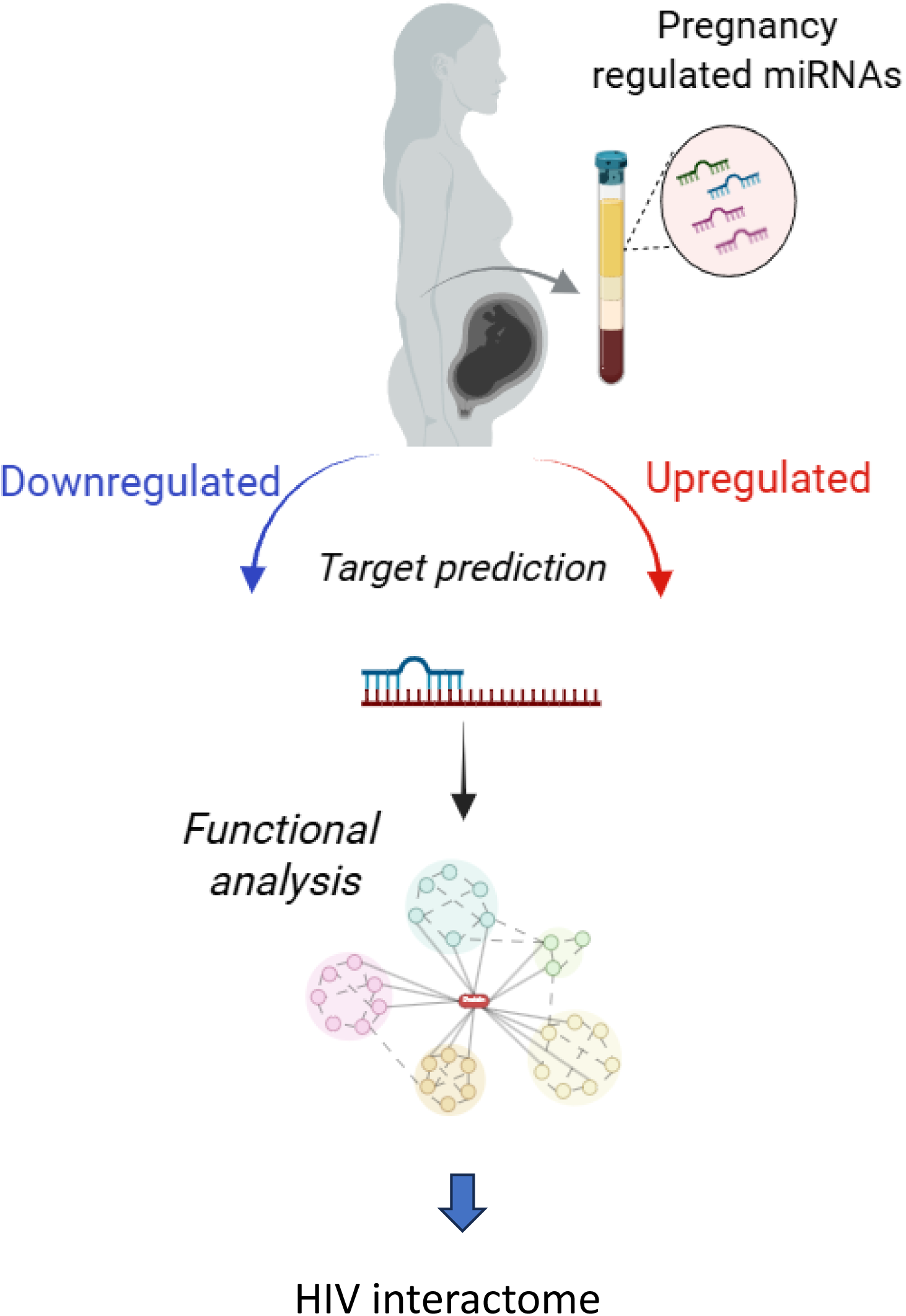

## 1 Introduction

Noncoding RNAs emerge as important regulators of maternal health (1). Among those miRNAs, which are post-transcriptional regulators of protein expression, and especially placental miRNAs, have been found to be associated with pregnancy complications, inflammation and fetal tolerance (2). While more rarely studied in pregnancy, individual miRNAs have been associated with viral infections, including HIV (3) and more recently miRNAs in the systemic circulation have been associated with the regulation of key drug metabolizing enzymes and transporter (BIORXIV/2025/665636). Pregnancy and the early postpartum period have been associated with higher risk of HIV(4); however, while hormonal pathways are suspected, underlying molecular mechanisms have not been elucidated. This study took the global transcriptome approach coupled with a robust bioinformatics pathway analysis to investigate the role of miRNAs differentially expressed during pregnancy and lactation in the risk of HIV acquisition through predicted enrichment of the HIV interactome defines as all host genes with know and experimentally validated role in HIV pathogenesis.

## 2 Methods

### 2.1 Ethical statement

The study received ethical approval from Brigham and Women’s Hospital (Boston, Massachusetts, USA). Institutional authorities in Uganda and Zimbabwe also approved the transfer of samples to the United States. Prior to enrollment, all participants provided informed consent after reading and understanding the study details. All methods and procedures adhered to the guidelines approved by the respective institutions.

### 2.2 Study population

Participants of this study were part of a larger multicentric cohort on hormonal contraception and HIV acquisition conducted in Uganda and Zimbabwe (5). A total of 88 HIV-negative women were enrolled, providing 174 serum samples in total. Among these, one participant contributed serum samples from two consecutive pregnancies, and 12 unpaired samples were included (1 pre-pregnancy and 11 during pregnancy). Women were tested for *Candida albicans* and a panel of sexually transmitted infections (STIs) consisting of human immunodeficiency virus (HIV), herpes simplex virus (HSV), *Chlamydia trachomatis*, *Neisseria gonorrhoeae* and *Trichomonas vaginalis* (6, 7). The presence of bacterial vaginosis (BV) was assessed at the study by experienced physicians, using the Nugent score (6). Table 1 summarizes population characteristics and clinical variables.

**Table 1.** Characteristics of the participating subjects by study site and clinical variables.

|  |  |  |  |
| --- | --- | --- | --- |
| **Variables** | **Uganda** | **Zimbabwe** | **Total (N)** |
| ***Study Groups: n (%)*** |  |  |  |
| Pre-pregnant | 18 (34.6%) | 34 (65.4%) | 52 |
| Pregnant | 39 (44.3%) | 49 (55.7%) | 88 |
| Breastfeeding | 15 (44.1%) | 19 (55.9%) | 34 |
| ***Age at first visit (years)*** |  |  |  |
| Mean ± SD | 23.05 ± 3.5 | 24.02 ± 4.2 | NA |
| ***Paired Samples: n*** |  |  |  |
| Pregnant – pre-pregnant | 13 | 30 | 43 |
| Pregnant – breastfeeding | 13 | 15 | 28 |
| ***STI Prevalence (Baseline): n (%)*** |  |  |  |
| HIV seroconversion <sup>1</sup> | 0 (0%) | 5 (100%) | 5 |
| Chlamydia | 1 (33.3%) | 2 (66.7%) | 3 |
| Gonorrhea | 3 (75%) | 1 (25%) | 4 |
| HSV | 31 (46.3%) | 36 (53.7%) | 67 |
| Trichomoniasis | 1 (%) | 2 (%) | 3 |
| ***Other vaginal infections*** |  |  |  |
| Candidiasis | 6 (30%) | 14 (70%) | 20 |
| **Bacterial vaginosis (Nugent score)** |  |  |  |
| BV (7 – 10) | 15 (38.5%) | 24 (61.5%) | 39 |
| Intermediate (4 – 6) | 9 (37.5%) | 15 (62.5%) | 24 |
| Normal (0 – 3) | 28 (45.2%) | 34 (54.8%) | 62 |
| Abbreviations: STI= sexually transmitted infections, HIV= human immunodeficiency virus, HSV= herpes simplex virus, BV= bacterial vaginosis, NA= not applicable. <sup>1</sup> Five women serum converted during the study in the groups pregnant (n=3) and breastfeeding (n=2). |  |  |  |

### 2.3 Whole human miRNA transcriptome profiling

The miRNA whole transcriptome (miRWT) from serum samples of women in pre-pregnancy, pregnancy, and breastfeeding stages was analyzed using the high-fidelity HTG EdgeSeq platform (HTG Molecular Diagnostics Inc., Tucson, AZ, USA). This platform uses a quantitative nuclease protection assay combined with PCR to attach adapters and barcodes, enabling an RNA extraction-free procedure. Following sample cleanup, a library pool was generated and amplified for next-generation sequencing (NGS) using the Illumina NextSeq instrument and the NextSeq 500/550 High Output Kit v2.5 (75 cycles). Samples were processed across three batches of miRNA transcriptome assay, which included 2083 mature miRNAs, 13 housekeeping genes a positive control (Supplemental Table 1) and 5 negative control genes (from *Arabidopsis thaliana aintegumenta*: ANT1, ANT2, ANT3, ANT4 and ANT5) (8).

### 2.4 Differential miRNA expression and statistical analysis

Low-expressing miRNAs were filtered out by retaining those with counts per million (CPM) > 4 in at least 10 samples, rendering 2,079 miRNAs. Voom transformation (9) was applied to convert counts to log-CPM, accounting for mean-variance relationships and assigning precision weights based on read depth. The log-CPM values were normalized using quantile normalization (10). Principal component analysis (PCA) was performed using the R package stats (v4.3.1) to assess the similarity and dissimilarity among samples based on their miRNA expression profiles. Differentially expressed miRNAs (DE-miR) between groups (pre-pregnancy, pregnancy, and breastfeeding) were identified using mixed-effect linear models using *dream* function (R package variancePartition v.1.30.2) (11), and tested for fixed-(group, country, plate, Nugent score, candidiasis and STIs) and random-effect (participant ID) variables. Missing values for candidiasis, Nugent score, and *T. vaginalis* were imputed due to inconsistencies between visits. If a value was missing at a later visit but available at the first visit, it was imputed using the previous visit’s data. If the first visit had missing data, the value was imputed using data from the subsequent visit. We calculated moderated t-statistics for differential expression using the *eBayes* function, which applies empirical Bayes moderation to shrink standard errors toward a global mean. Volcano plots label top miRNAs by smallest *p*-values or largest log-FC. Heatmaps plot clipped z-scores of adjusted log-CPM for top miRNAs selected by *p*-values across comparisons.

### 2.5 Construction of a HIV-interactome dataset

We created a HIV-interactome gene dataset by combining human genes identified in a previous study (12) with those from the manually curated ‘HIV-1 Human Interaction Database’ (13). The database was maintained by The National Center for Biotechnology Information (NCBI) until June 2024 and downloaded by us on 02/10/2023. It compiles reports on HIV-human protein-protein interactions (“protein interactions”) and HIV replication and infectivity (“replication interactions”). The Luo et al. report comprises 237 genes, while the ‘HIV-1 Human Interaction Database’ contains 4,577 genes, with an overlap of 211 genes. Thus, our HIV-interactome dataset includes 4,603 genes for analysis (Supplemental Table 2). The HIV-interactome genes were annotated using SYNGO ID conversion tool (14).

### 2.6 Identification of miRNA-Targets

DE-miR (FDR < 0.1) in the comparisons between Pregnant vs. Pre-pregnant and Pregnant vs. Breastfeeding groups were overlapped to identify miRNAs up- and downregulated during pregnancy. The miRNAs (miRBase V22) were queried in the miRWalk database (release_2022_01) (15) to identify empirically validated target genes. The identified targets were annotated (14) and then utilized for Modular Enrichment Analysis (MEA). All comparative set analyses were conducted using Venn diagrams generated with InteractiVenn (15), while the UpSet plot (16) was created using the UpSetR package (V1.4.0) (17).

### 2.7 Modular Enrichment Analysis

All the validated predicted targets of DE-miR in pregnancy were subjected to Modular Enrichment Analysis (MEA) by applying ClueGo v2.5.9 (16) and CluePedia v1.5.9 (3), implemented as Cytoscape (v3.10.3) applications. The MEA identifies biologically meaningful patterns by analyzing groups (modules) of related genes or proteins. It employs *Kappa* statistics and *p*-value calculations to assess enrichment, accounting for the redundant and networked nature of biological annotations (17). The upregulated (Cluster 1) and downregulated (Cluster 2) gene sets were loaded into ClueGo. Non-redundant biological information was obtained from Gene Ontology of Immune System (GO-IS) and Kyoto Encyclopedia of Genes and Genomes (KEGG) by merging redundant groups with ≥ 50% gene overlap and preserving the more representative terms per their association strength (*Kappa* score ≥ 0.6). The statistical significance of enriched terms was estimated by enrichment/depletion two-sided hypergeometric test and corrected by the Bonferroni step-down test (p-value cutoff ≤ 0.05). HIV-interactome genes were mapped using the AutoAnnotate app (v1.5.2) (18), and hub genes were identified through topological analysis of the PPI networks using the cytoHubba (v0.1) (19) as described (8). Protein–protein interaction (PPI) and hub genes-targeting miRNAs networks were visualized with the Organic Layout algorithm (yFiles Layout Algorithms, v1.1.4). Predicted interactions between HIV-interactome, hub genes and their associated enriched pathways were uploaded to Cytoscape, and the network was visualized using Degree Sorted Circle Layout, followed by the generation of a hub elements subnetwork using cytoHubba. The subnetwork was visualized using Organic Layout.

## 3 Results

### 3.1 Serum miRNA Expression Changes Across Stages of Female Reproductive Status in Longitudinal Samples

The PCA plots comparing sample variance showed that the samples of pregnant (FIG 1A) and breastfeeding (Fig. 1B) women were globally different from those of preconception women. The heatmaps illustrate the top 50 DE miRNAs across samples for the Pregnancy vs. Pre-pregnancy (Figure 1C) and Breastfeeding vs. Pregnant (Figure 1E) comparisons. Pregnancy vs. Pre-pregnant comparison revealed 684 DE miRNAs and the Pregnancy vs. Breastfeeding revealed 240 DE miRNAs with FDR<0.1) (Supplemental Table 3), corresponding to 32.8% of the miRWT, and 11.5% of the miRWT, respectively. Volcano plots annotate the miRNAs with lowest FDR and largest fold change in both comparisons (Fig. 1 D and F)

**Figure 1:**
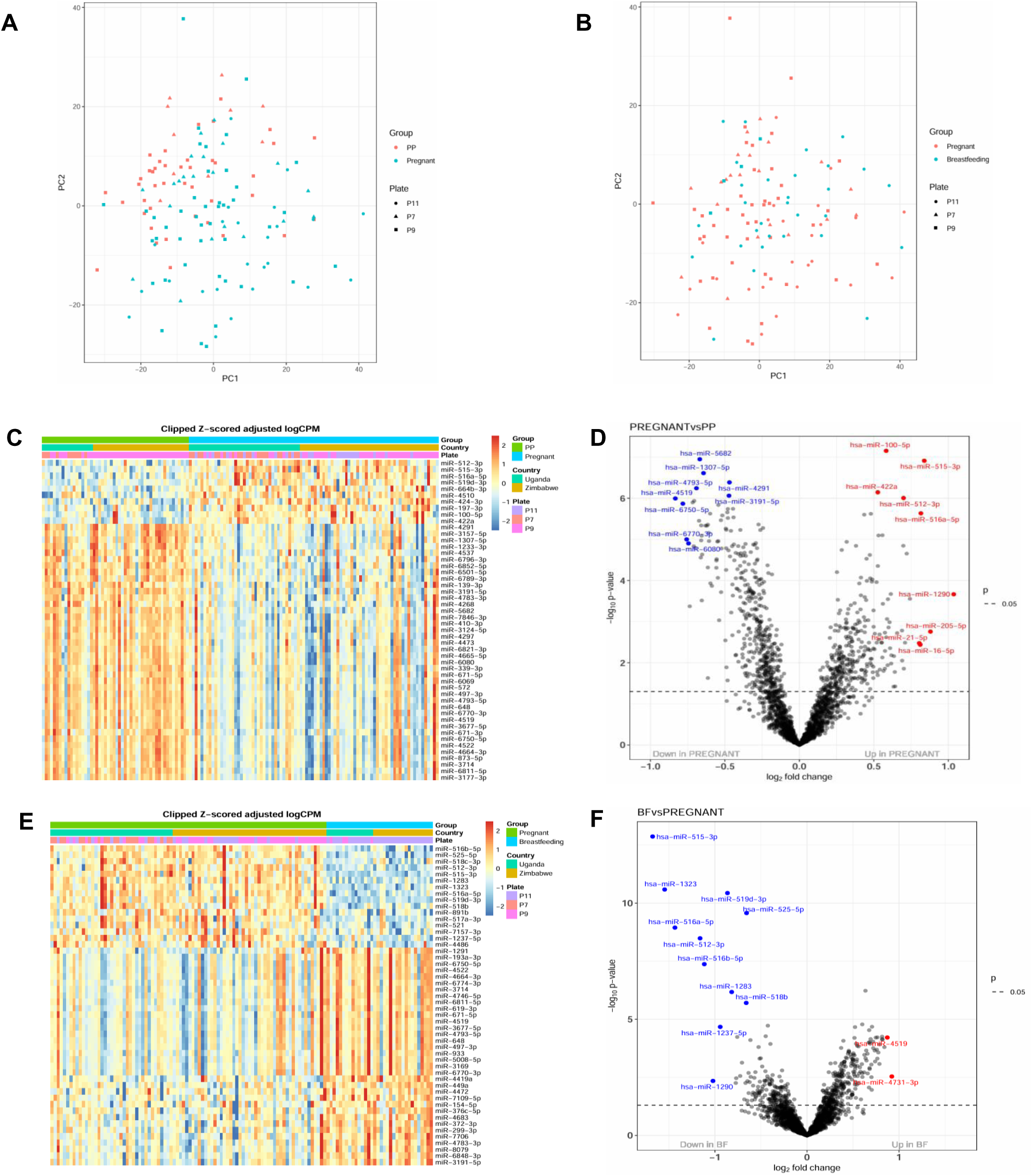
Serum miRNA expression by female reproductive status. (**A, B**) Principal Component Analysis (PCA) was performed for (**A**) Pregnant (P) vs. Pre-pregnant (PP) and (**B**) Breastfeeding (BF) vs. Pregnant groups, including batch effect (Plate). **(C, E**) Heatmaps display the top 50 DE miRNAs (rows) across individual samples (column) for the comparisons P vs. PP (**C**) and BF vs. P (**E**) with clustering by country and batch effect (Plate). Expression levels are represented by color intensity: red for upregulation and blue for downregulation. (**D, F**) Volcano plots show differentially expressed (DE) miRNAs for P vs. PP (**D**) and BF vs. P (**F**). The most significantly downregulated (blue) and upregulated (red) miRNAs based on smallest p-values or largest log fold changes (log FC) are annotated.

### 3.2 Pregnancy-Driven Serum miRNAs Target Genes Shared with the HIV Interactome

Pregnancy-driven serum miRNAs identified by the overlap between the Pregnant vs. Prepregnant and Pregnant vs. Breastfeeding comparisons included 160 DE-miRNAs (FDR <0.1), of which 29 were upregulated and 131 downregulated (Supplemental Table 4). These DE-miRNAs were predicted to, respectively, downregulate 644 genes and upregulate 2444 genes (Supplemental Table 5). Among them, 98 (15.2%) of downregulated genes and 629 (25.7%) of upregulated genes overlapped with the HIV interactome dataset (Figure 2A), while 119 proteins fell into the intersection of the 3 groups and 216 were predicted to be bidirectionally expressed (both up and downregulated) (Figure 2A, Supplemental Table 6).

**Figure 2:**
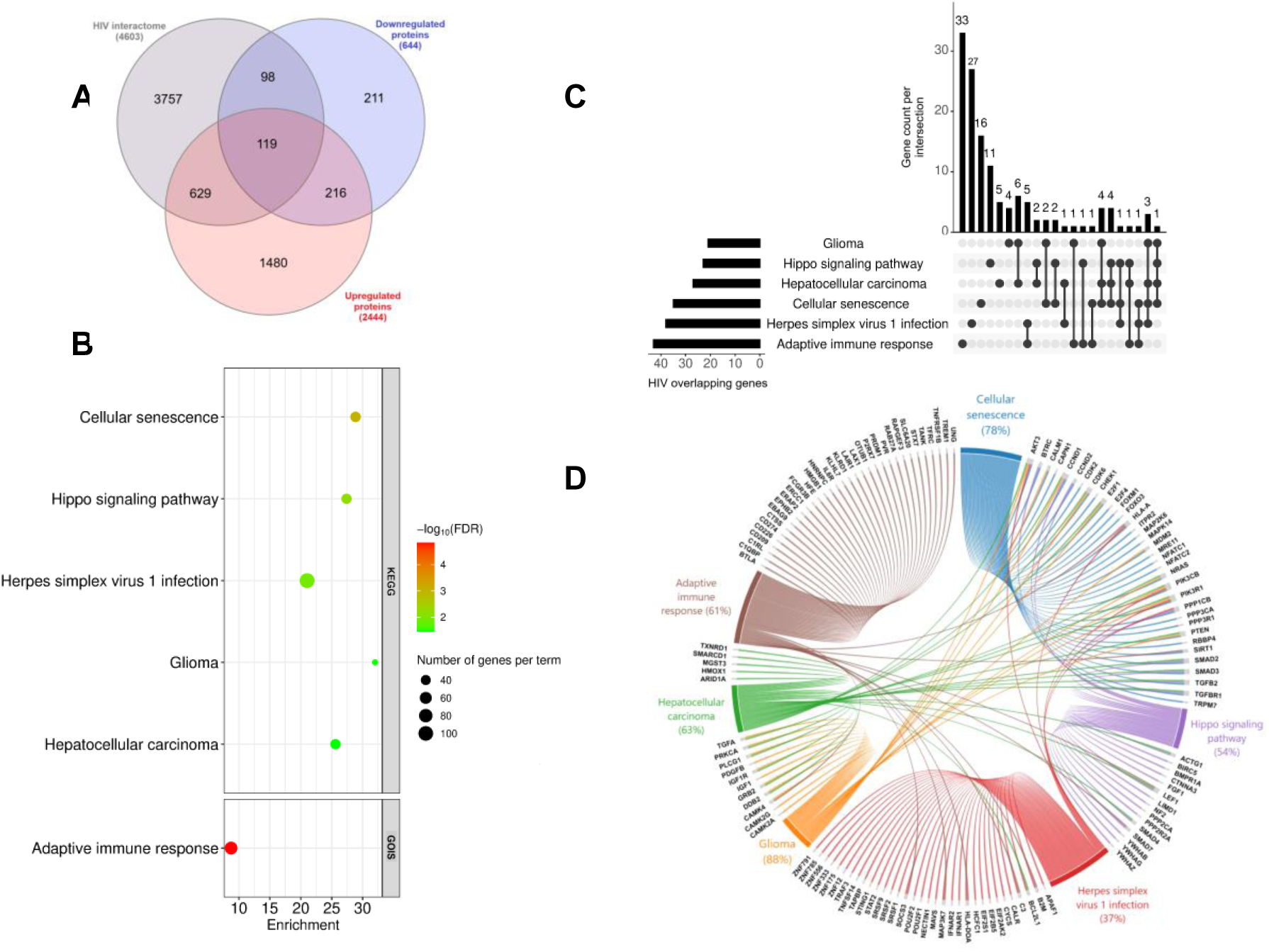
Modular enrichment analysis (MEA) of validated targets of serum miRNAs altered during pregnancy and enriched in the overlap with the HIV interactome. The Venn diagram (**A**) shows the overlap (intersections) and unique elements (differences) between the HIV interactome and the validated targets of pregnancy-driven serum miRNAs. MEA was conducted using all validated targets (2444 upregulated and 644 downregulated) of pregnancy-driven serum miRNAs and presented as Bubble plot (**B**). The Y-axis of the bubble plot annotates the names of enriched pathways (KEGG, upper plot) and terms (GOIS, lower plot) and the X-axis shows enrichment as % of genes per pathway or term. Bubble color represents the significance level of each pathway/term (-log10 FDR), while size indicates the total number of genes enriched in each pathway/term. The UpSet plot (**C**) and the cord plot (**D**) highlight the HIV interactome gene overlap contributing to the enrichment of the biological pathways shown in (B). The Y-axis of upper graph on the UpSet plot shows the count of HIV overlapping genes that are either unique to or shared among specific pathway combinations, as indicated by the connected dots below the X-axis. Horizontal bars on the lower left of the Upset plot represent the total number of HIV-overlapping genes associated with each individual enriched pathway. The chord plot illustrates all genes within enriched biological terms that overlap with the HIV interactome and their distribution across pathways. Ribbons and colors in the cord plot distinguish the 6 enriched biological groups/pathways. The percentages below the labels indicate the proportion of genes in each term that are shared with the HIV interactome.

### 3.3 The Genes Targeted by Pregnancy-Driven Serum miRNAs are Linked to Adaptive Immune Response, Hippo Signaling, Cancer, Cellular Senescence and Viral Infection

All confirmed targets of pregnancy-modulated serum miRNAs were employed to identify associated biological pathways. Targets were found to be functionally associated with ‘Adaptive Immune Response’ (GO:0002250), ‘Cellular Senescence’ (KEGG:04218), ‘Hippo Signaling Pathway’ (KEGG:04390), ‘Herpes Simplex Virus 1 (HSV1) Infection’ (KEGG:05168), ‘Glioma’ (KEGG:05214) and ‘Hepatocellular Carcinoma’ (HCC) (KEGG:05225) with an FDR <0.05 (Figure 2B). The GOIS term ‘Adaptive Immune Response’ was the most significant (corrected *p* = 0.000015) while ‘Cellular Senescence’ showed the greatest enrichment score (28.85% of genes per pathway) (Figure 2B, Supplemental Table 7). Over 71% of the genes associated with those 6 biological functions are predicted to be upregulated as a result of downregulated pregnancy-driven DE miRNAs (Supplemental Table 7). Among the genes contributing to enriched biological pathways, 132 were found to overlap with the HIV interactome (Figure 2C, Supplemental Table 8), highlighting shared molecular mechanisms. Of these, 36 genes are shared across the 6 key pathways identified (Figure 2C-D, Supplemental Table 8).

Compared to the ‘Glioma’, ‘Cellular Senescence’, ‘Hippo Signaling Pathways’ and ‘HCC’, those related to viral infection and immune response contained a higher number of HIV-interactome genes exclusively driving their enrichment, i.e. genes not shared with the other pathways (Figure 2C). In this sense, the ‘HSV1 infection’ pathway and Adaptive Immune Response term included 27 and 33 exclusive HIV-interactome genes, respectively (Figure 2C, Supplemental Table 8). Functional connections between enriched biological pathways and HIV-overlapping genes targeted by pregnancy-driven serum miRNAs (Figure 2D) show that cancer-related pathways and the ‘Cellular Senescence’ function share the largest number of HIV-overlapping proteins. The percentage of genes shared with the HIV interactome in each pathway ranged from 37% to 88%.

### 3.4 PPI Network Mapping Reveals Hub Gene Interactions and miRNA-Hub Gene networks Highlight Pregnancy-Driven miRNA Control of Hub Genes

We constructed PPI networks for genes targeted by pregnancy-driven miRNAs shared with the HIV interactome. These genes contributed to six enriched biological pathways and were functionally mapped. ‘Adaptive Immune Response’ PPI network (Figure 3A) included 43 nodes and 80 edges mapped into 2 functional groups (*cellular receptors* and *DNA processes*), while ‘Cellular Senescence’ comprised 35 nodes and 180 edges assigned to 5 functional groups (*cell cycle, calcium-activated phosphatase, ion channel, MAPK signaling and plasma membrane proteins*) (Figure 4A). ‘Hippo Signaling Pathway’ included 23 nodes and 69 edges distributed into 3 functional groups (*TGF-β signaling, cellular housekeeping and protein phosphatases*) (Figure 5A). The ‘HSV-1 Infection’ PPI network (Figure 6A) included 38 nodes and 77 edges mapped into 5 functional groups (*IFN signaling, antigen presentation and complement pathway, cellular processes, PI3K/AKT signaling pathway* and *serine/arginine-rich (SR) protein family*). The cancer related pathways, ‘Glioma’ and ‘HCC’ included 21 nodes and 102 edges associated with *cell protein kinase* (Figure 7A), and 27 nodes and 131 edges linked to *Cell Growth and DNA repair, respectively* (Figure 8A). In sum, network mapping revealed essential functional groups likely altered in pregnancy.

**Figure 3.**
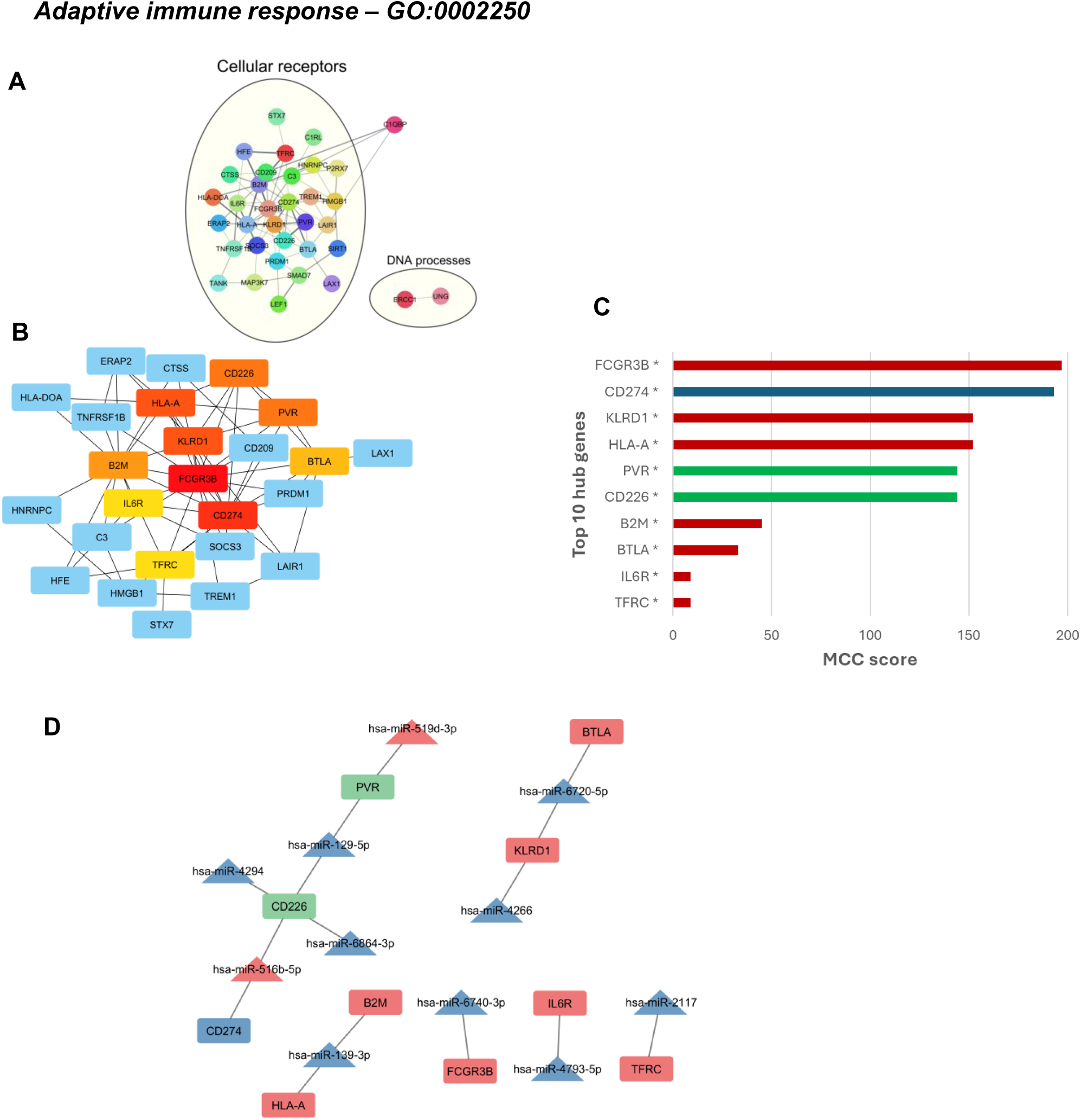
Protein-protein interaction (PPI) and hub gene networks of HIV-interactome genes within the Adaptive Immune Response GO term, including predicted miRNA–target interactions. (**A**) protein-protein interaction (PPI) networks display HIV-interactome genes contributing to the adaptive immune response term enrichment. Functional annotations highlight commonalities in protein roles. **(B**) PPI network of the top 10 hub genes within the term, identified using cytoHubba based on Maximal Clique Centrality (MCC) analysis. Node colors indicate MCC scores, from high (red) to low (yellow). (**C**) Bar plots of the top 10 hub genes by MCC score. Asterisks indicate genes annotated under cellular receptor-related functions, as shown in Figures 3A. Bar colors represent the predicted regulation of each gene in pregnancy based on differential miRNA expression: blue for downregulated, red for upregulated, and green for mixed regulation (both up- and downregulated). (**D**) miRNA–hub gene target networks show 21 nodes and 15 edges. Triangles represent miRNAs and rectangles represent hub genes. Node colors reflect inferred regulation in pregnancy: blue (downregulated), red (upregulated), and green (mixed).

**Figure 4:**
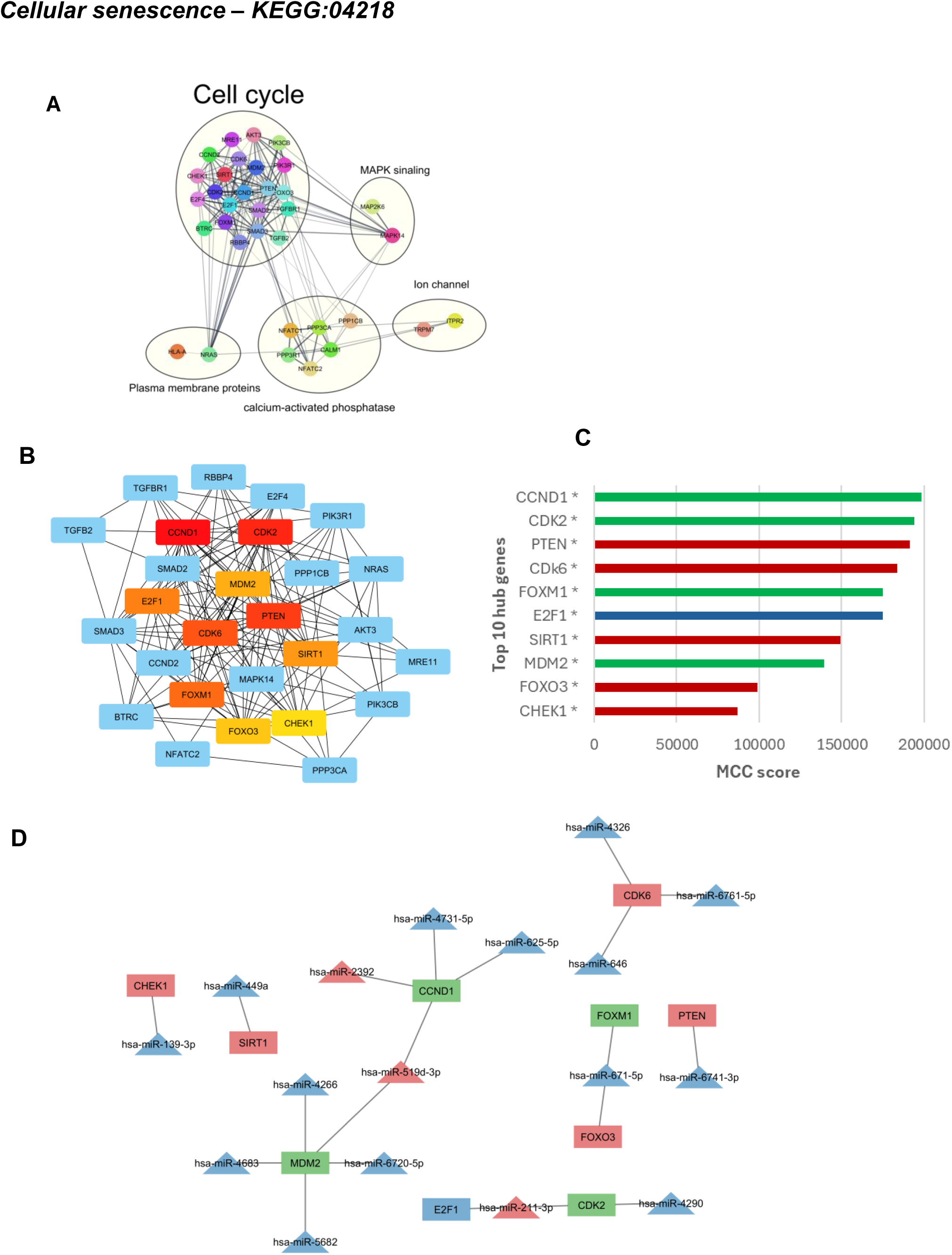
PPI and hub gene networks of HIV-interactome genes within the Cellular Senescence KEGG pathway, including predicted miRNA–target interactions. (**A**) protein-protein interaction (PPI) networks display HIV-interactome genes contributing to the cellular Senescence pathway enrichment. Functional annotations highlight commonalities in protein roles. (**B**) PPI network of the top 10 hub genes within the term, identified using cytoHubba based on Maximal Clique Centrality (MCC) analysis. Node colors indicate MCC scores, from high (red) to low (yellow). (**C**) Bar plots of the top 10 hub genes by MCC score. Asterisks indicate genes annotated under in cell cycle, as shown in Figures 4A. Bar colors represent the predicted regulation of each gene in pregnancy based on differential miRNA expression: blue for downregulated, red for upregulated, and green for mixed regulation (both up and downregulated). (**D**) miRNA–hub gene target networks show 27 nodes and 20 edges. Triangles represent miRNAs and rectangles represent hub genes. Node colors reflect inferred regulation in pregnancy: blue (downregulated), red (upregulated), and green (mixed).

**Figure 5:**
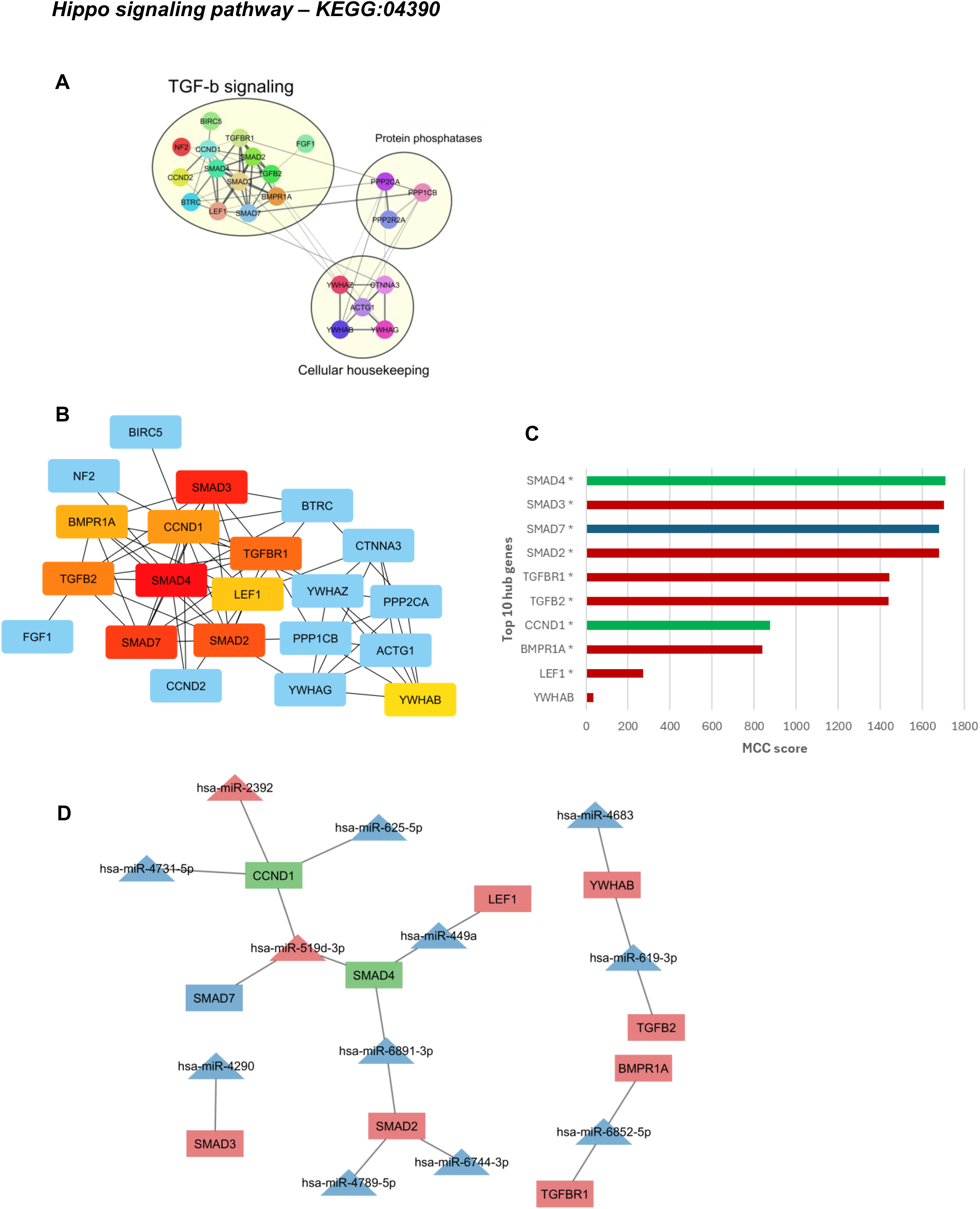
PPI and hub gene networks of HIV-interactome genes within the Hippo Signaling KEGG pathway, including predicted miRNA–target interactions. (**A**) protein-protein interaction (PPI) networks display HIV-interactome genes contributing to the Hippo Signaling pathway enrichment. Functional annotations highlight commonalities in protein roles. (**B**) PPI network of the top 10 hub genes within the term, identified using cytoHubba based on Maximal Clique Centrality (MCC) analysis. Node colors indicate MCC scores, from high (red) to low (yellow). (**C**) Bar plots of the top 10 hub genes by MCC score. Asterisks indicate genes annotated under TGFβsignaling, as shown in Figures 5A. Bar colors represent the predicted regulation of each gene in pregnancy based on differential miRNA expression: blue for downregulated, red for upregulated, and green for mixed regulation (both up and downregulated). (**D**) miRNA–hub gene target networks show 22 nodes and 18 edges. Triangles represent miRNAs and rectangles represent hub genes. Node colors reflect inferred regulation in pregnancy: blue (downregulated), red (upregulated), and green (mixed).

**Figure 6:**
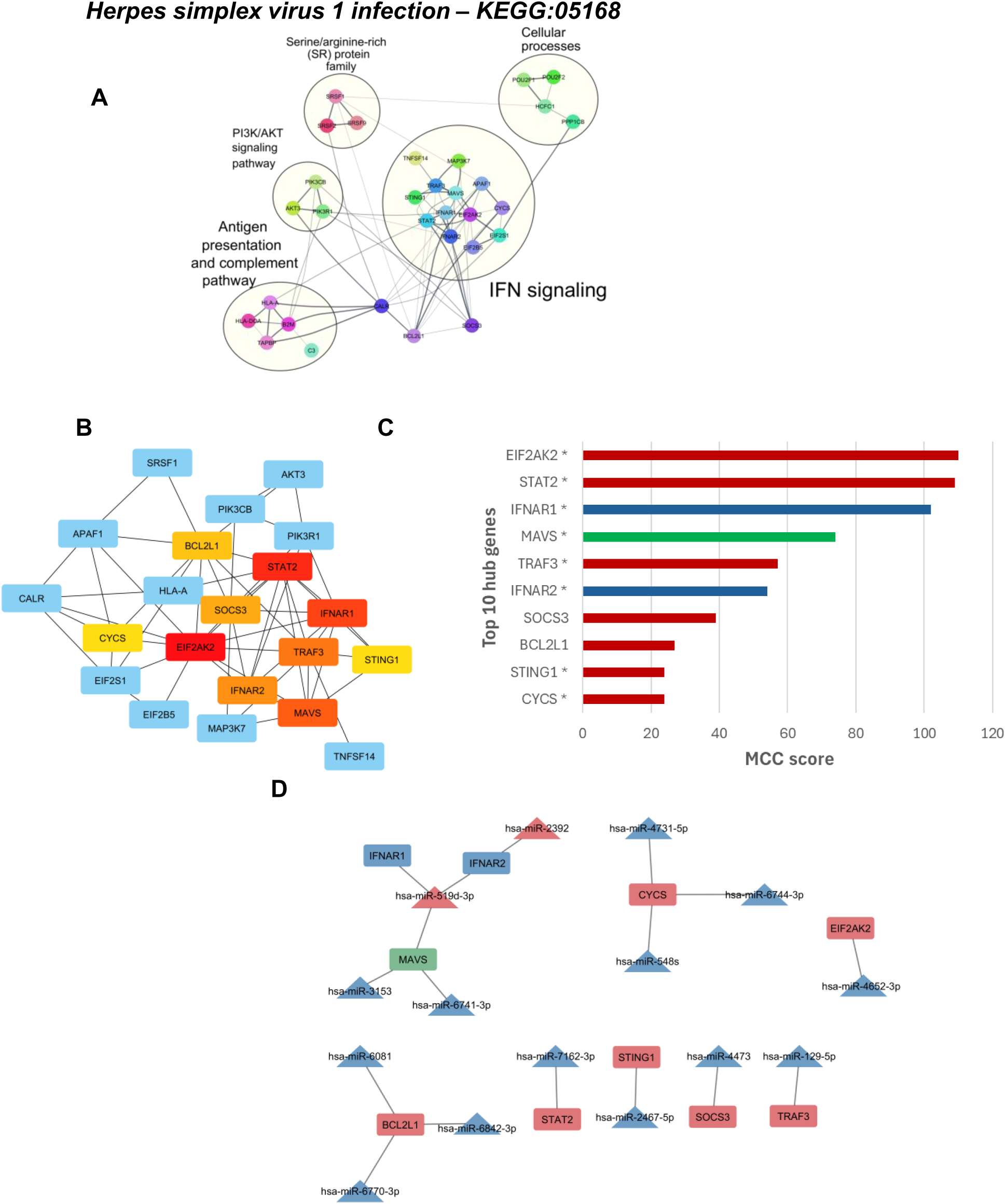
PPI and hub gene networks of HIV-interactome genes within the Herpes Simplex Virus 1 Infection KEGG pathway, including predicted miRNA–target interactions. **(A**) protein-protein interaction (PPI) networks display HIV-interactome genes contributing to the Herpes Simples Virus 1 (HSV1) infection pathway enrichment. Functional annotations highlight commonalities in protein roles. (**B**) PPI network of the top 10 hub genes within the term, identified using cytoHubba based on Maximal Clique Centrality (MCC) analysis. Node colors indicate MCC scores, from high (red) to low (yellow). (**C**) Bar plots of the top 10 hub genes by MCC score. Asterisks indicate genes annotated under IFN signaling, as shown in Figures 6A. Bar colors represent the predicted regulation of each gene in pregnancy based on differential miRNA expression: blue for downregulated, red for upregulated, and green for mixed regulation (both up- and downregulated). (**D**) miRNA–hub gene target networks show 25 nodes and 17 edges. Triangles represent miRNAs and rectangles represent hub genes. Node colors reflect inferred regulation in pregnancy: blue (downregulated), red (upregulated), and green (mixed).

**Figure 7:**
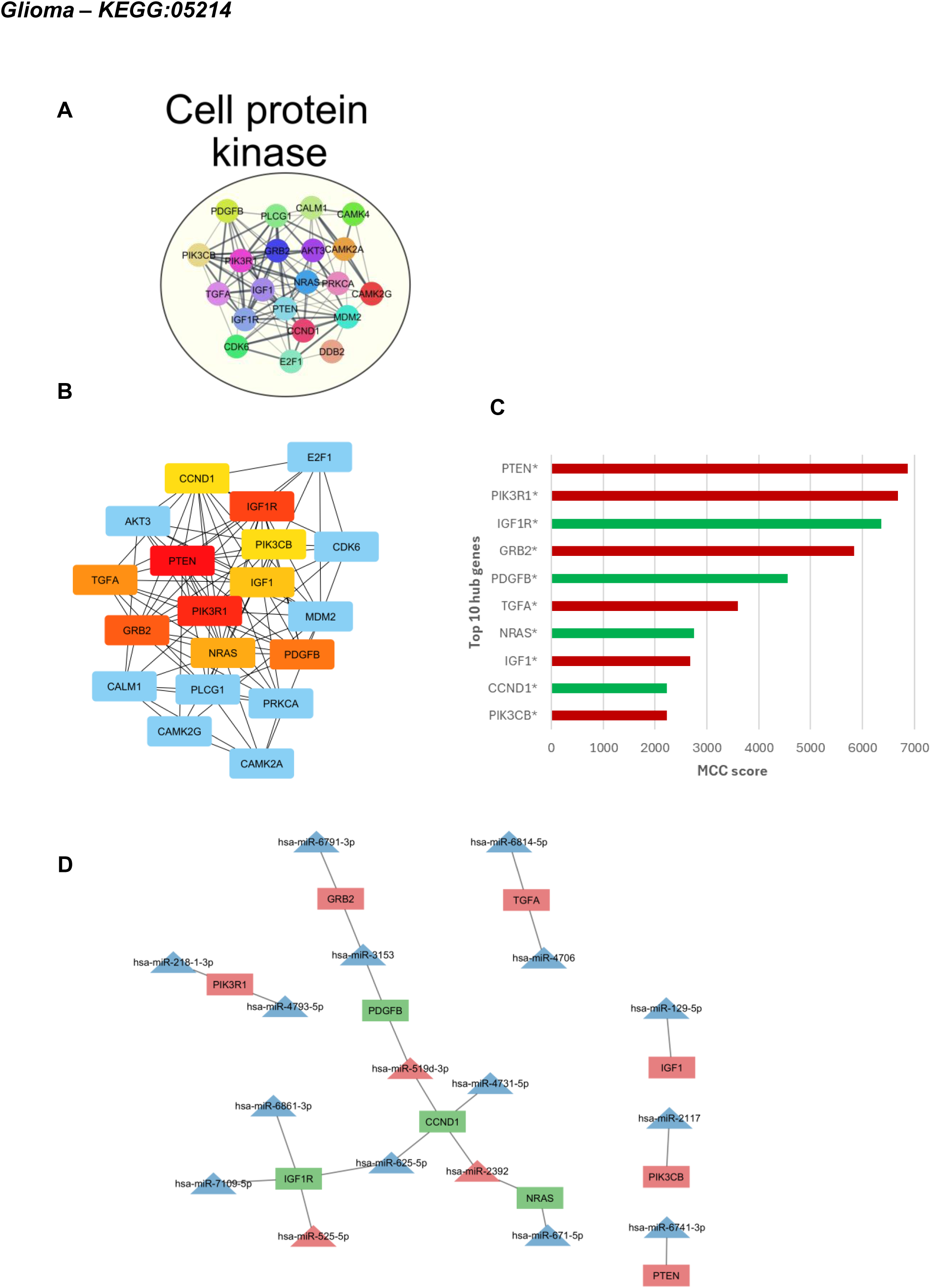
PPI and hub gene networks of HIV interactome genes within the Glioma KEGG pathway, including predicted miRNA–target interactions. (**A**) protein-protein interaction (PPI) networks display HIV interactome genes contributing to the Glioma pathway enrichment. Functional annotations highlight commonalities in protein roles. (**B**) PPI network of the top 10 hub genes within the term, identified using cytoHubba based on Maximal Clique Centrality (MCC) analysis. Node colors indicate MCC scores, from high (red) to low (yellow). (**C**) Bar plots of the top 10 hub genes by MCC score. Asterisks indicate genes annotated under Cell protein kinase, as shown in Figures 7A. Bar colors represent the predicted regulation of each gene in pregnancy based on differential miRNA expression: blue for downregulated, red for upregulated, and green for mixed regulation (both up and downregulated). (**D**) miRNA–hub gene target networks show 27 nodes and 21 edges. Triangles represent miRNAs and rectangles represent hub genes. Node colors reflect inferred regulation in pregnancy: blue (downregulated), red (upregulated), and green (mixed).

**Figure 8:**
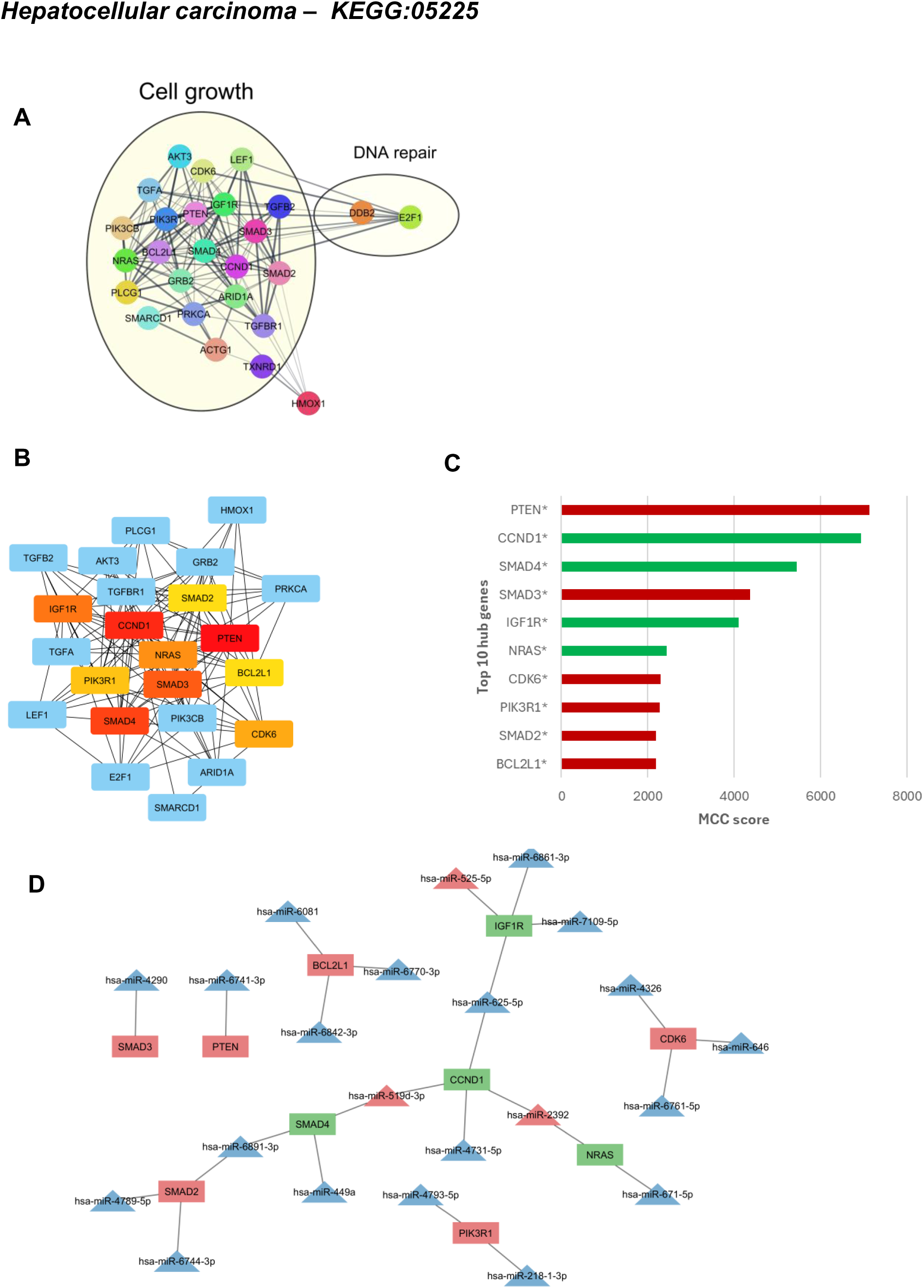
PPI and hub gene networks of HIV interactome genes within the Hepatocellular Carcinoma (HCC) KEGG pathway, including predicted miRNA–target interactions. (**A**) protein-protein interaction (PPI) networks display HIV interactome genes contributing to the HCC pathway enrichment. Functional annotations highlight commonalities in protein roles. (**B**) PPI network of the top 10 hub genes within the term, identified using cytoHubba based on Maximal Clique Centrality (MCC) analysis. Node colors indicate MCC scores, from high (red) to low (yellow). (**C**) Bar plots of the top 10 hub genes by MCC score. Asterisks indicate genes annotated under Cell growth, as shown in Figures 8A. Bar colors represent the predicted regulation of each gene in pregnancy based on differential miRNA expression: blue for downregulated, red for upregulated, and green for mixed regulation (both up and downregulated). (**D**) miRNA–hub gene target networks show 32 nodes and 26 edges. Triangles represent miRNAs and rectangles represent hub genes. Node colors reflect inferred regulation in pregnancy: blue (downregulated), red (upregulated), and green (mixed).

Hub genes are highly connected within a PPI network. They are prime candidates for drug targeting, as their central roles within the network suggest they are likely essential for disease initiation and progression. To further explore this, we identified the top 10 hub genes that support and maintain the structural integrity of the all the PPI networks (Figure 3B–8B), presenting them according to their MCC scores and predicted direction of regulation, either upregulated, downregulated or bidirectional (Figure 3C–8C). We identified pregnancy-driven miRNAs that regulate the hub genes and constructed their corresponding regulatory networks, highlighting the predicted miRNA-hub gene interactions within the following pathways: ‘Adaptive Immune Response’ (Figure 3D), ‘Cellular Senescence’ (Figure 4D), ‘Hippo Signaling Pathway’ (Figure 5D), ‘HSV1 Infection’ (Figure 6D), ‘Glioma’ (Figure 7D), and ‘HCC’ (Figure 8D). Our data show a total of 47 unique hub genes across all the enriched pathways (Table 2) likely under post-transcriptional “one-to-many” and “many-to-one” regulation by 45 pregnancy-driven miRNAs, of which 40 are downregulated and 5 are upregulated.

**Table 2.**
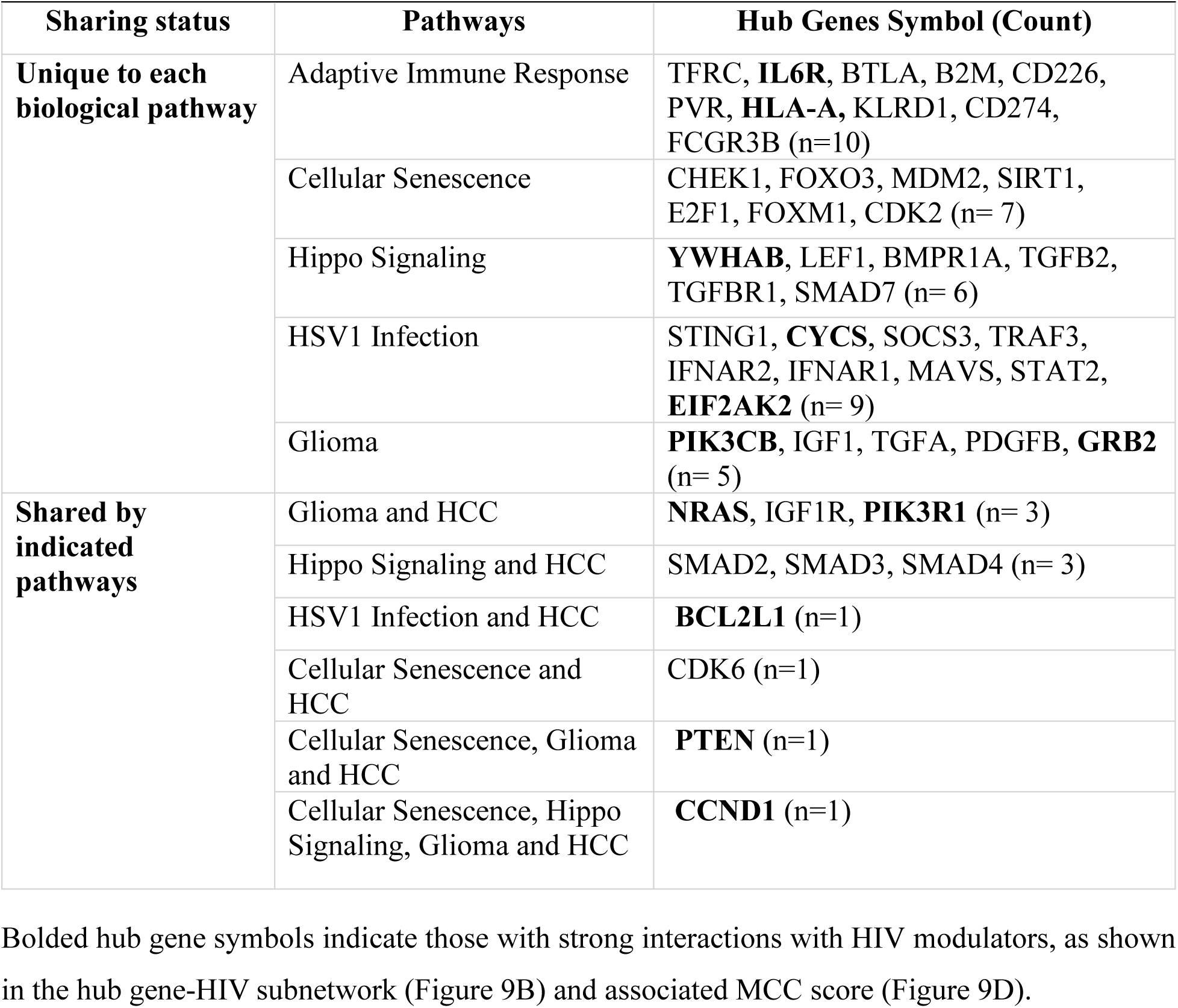
Comparison of Hub Genes Shared Between or Exclusive to Pregnancy Enriched Pathways.

### 3.5 Hub Genes are Co-targeted by Pregnancy-Driven miRNAs and HIV modulators

Interactions between hub genes and HIV modulators were sourced from HIV-NCBI database (13), and compiled into tables (Table 3-7). These data were leveraged to construct a network incorporating the pathways associated with each hub gene (Figure 9A). We found that the hub genes were reported to interact with 19 HIV modulators, including an HIV biological process (HIV replication) and 18 HIV proteins (Capsid, gp120, gp160 – precursor of gp120 and gp41, gp41, Integrase, Matrix, Nef, Nucleocapsid, p6, Pol, Pr55(gag) – (gag polyprotein precursor, Retropepsin – a protease, Reverse transcriptase, Tat, Vif, Vpr and Vpu). Among these modulators, Tat and gp120 showed the highest connectivity within the network, followed by Nef, HIV-1 virus replication, Vpr, gp160, and Vpu (Figure 9C). HLA-A stood out as the most highly connected host protein, with BCL2L1 ranking second, followed by EIF2AK2, PTEN, and CYCS, completing the top 5 host proteins (Figure 9D). Taking together, our study reveals a complex interplay between hub genes potentially co-targeted by pregnancy-associated host miRNAs and HIV modulators during pregnancy.

**Figure 9.**
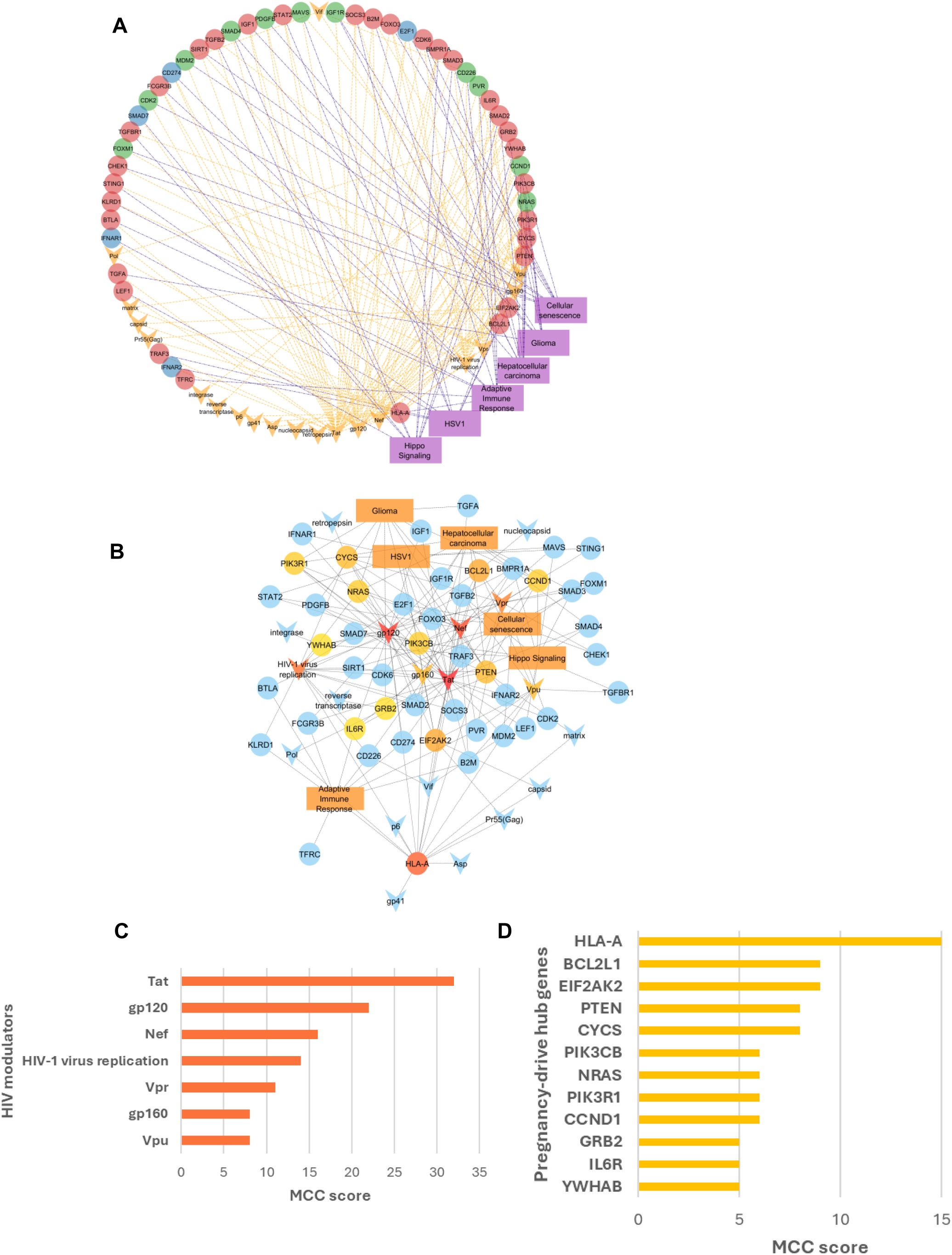
Circular network of hub genes, enriched pathways, and HIV-interacting modulators in pregnancy. (**A**) Ellipses represent hub genes with colors indicating their predicted regulation during pregnancy: downregulated (blue), upregulated (red), and mixed regulation (green). Enriched pathways are shown as purple rectangles, and HIV-interacting proteins are depicted as orange V-shapes. Orange dashed lines represent interactions between hub genes and HIV proteins, while purple dash-dot lines represent interactions between hub genes and enriched pathways. The network contains 72 nodes and 189 edges. (**B**) This subnetwork highlights the highly connected HIV proteins and miRNA targets – extracted from A – visualized using an orange color spectrum. The MCC score for HIV modulators (C) and pregnancy-driven hub genes (D) are shown. Pathways in pregnancy showed an MCC= 10.

**Table 3.** Interactions Between Adaptive Immune Response Hub Genes and HIV Proteins.

| Hub gene rank | Hub gene symbol | Hub gene name | HIV modulation and overall impact on hub gene |
| --- | --- | --- | --- |
| 1 | ↑FCGR3B * | Fc gamma receptor IIIb | ▪ *Envelope gp120* binds<br>▪ *Tat* inhibits and upregulates |
| 2 | ↓CD274/PDL1 * | CD274 molecule | ▪ *Envelope gp120* upregulates<br>▪ *Tat* upregulates |
| 3 | ↑HLA-A * | Major histocompatibility complex, class I, A | ▪ HLA-A is extensively targeted by HIV — *Nef* alone downregulates it in over 30 distinct interactions and also degrades, binds, and relocalizes it.<br>▪ *Tat*, *Vpu*, *Vpr*, and *gp120/gp160/gp41* contribute to either suppression or enhancement of HLA-A expression and activity. |
| 4 | ↑KLRD1 * | Killer cell lectin like receptor D1 | ▪ *HIV1 replication* upregulates expression |
| 5 | ↑↓CD226 * | CD226 molecule | ▪ *Nef* interacts<br>▪ *Vif* downregulates<br>▪ *HIV1 replication* downregulates |
| 6 | ↑↓PVR/CD155 * | PVR cell adhesion molecule | ▪ *Nef* downregulates, interacts, relocalizes<br>▪ *Vpu* downregulates, relocalizes<br>▪ *Vpr* upregulates |
| 7 | ↑B2M * | Beta2microglobulin | ▪ *Nef*, *Vpu*, and *Tat* downregulate |
| 8 | ↑BTLA * | B and T lymphocyte associated | ▪ *Envelope gp120* downregulates |
| 9 | ↑TFRC *§ | Transferrin receptor | ▪ *Unknown interaction (12) and not present in the HIVNCBI database* |
| 10 | ↑IL6R * | Interleukin 6 receptor | ▪ *gp120*, *gp160* upregulate<br>▪ *Tat* upregulates |

**Table 4.** Interactions Between Cellular Senescence Hub Genes and HIV Proteins.

| Hub gene rank | Hub gene symbol | Hub gene name | HIV modulation and overall impact on hub gene |
| --- | --- | --- | --- |
| 1 | ↑↓CCND1 *‡ | Cyclin D1 | ▪ Nef upregulates Cyclin D1; Tat has context-dependent effects (both up and downregulation) on Cyclin D1 expression |
| 2 | ↑↓ CDK2 * | Cyclin dependent kinase 2 | ▪ **Tat** stimulates CDK2/Cyclin E activity; phosphorylates<br>▪ Tat; regulates HIV transcription and host cell cycle progression |
| 3 | ↑PTEN *§ | Phosphatase and tensin homolog | ▪ **Tat** modulates PTEN expression and phosphorylation;<br>▪ **Nef** and **gp120** interact<br>▪ **matrix** proteins activate PTEN and AKT pathways |
| 4 | ↑CDK6 *† | Cyclin dependent kinase 6 | ▪ **Tat** reduces CDK6 abundance; HIV infection also downregulates CDK6 expression |
| 5 | ↑↓ FOXM1 * | Forkhead box M1 | ▪ **Vpr** downregulates FOXM1 expression |
| 6 | ↓E2F1 * | E2F transcription factor 1 | ▪ **gp120** activates E2F1 inducing neuronal death<br>▪ E2F1 inhibits **Tat** transcriptional activity |
| 7 | ↑SIRT1 * | Sirtuin 1 | ▪ **Tat** binds and inhibits SIRT1 deacetylase; downregulates via miRNAs; affects NFκB activation and HIV transcription |
| 8 | ↑↓ MDM2 * | MDM2 protooncogene | ▪ MDM2 ubiquitinates **Tat** (Lys71)<br>▪ **Vif** binds MDM2 and inhibits its activity<br>▪ MDM2 degrades **Vif**, controls TP53 stability |
| 9 | ↑FOXO3 * | Forkhead box O3 | ▪ **Tat** increases nuclear FOXO3a<br>▪ **Vpr** inhibits insulin/Akt-induced FOXO3 translocation<br>▪ FOXO3 supports **HIV replication** |
| 10 | ↑CHEK1 * | Checkpoint kinase 1 | ▪ **Vpr** activates ATR → phosphorylates Chk1 (Ser345) → ATR-mediated DNA damage response → G2/M arrest |

**Table 5.**
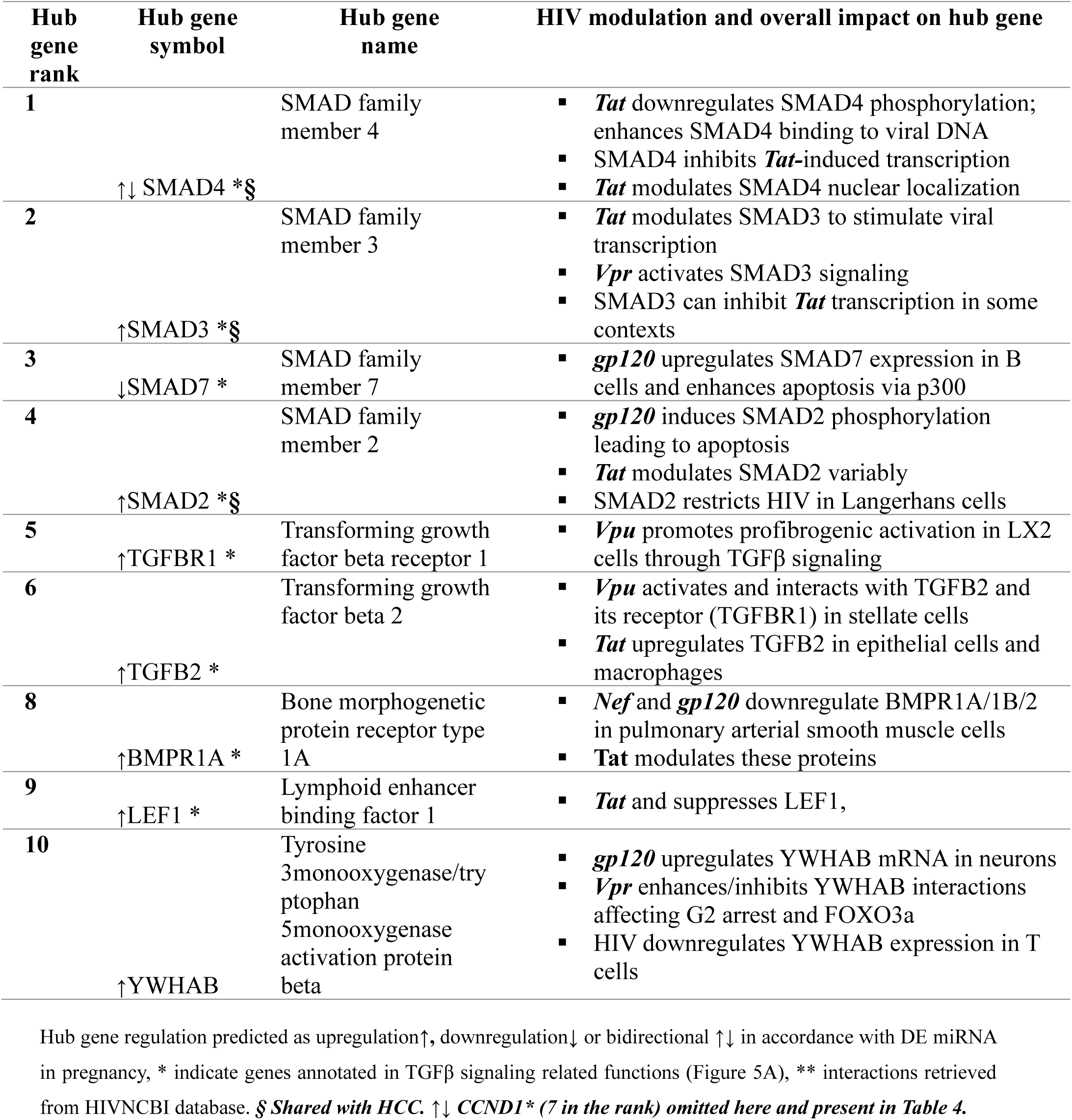
Interactions Between Hippo Signaling Pathway Hub Genes and HIV Proteins.

**Table 6.** Interactions Between HSV1 Infection Hub Genes and HIV Proteins.

| Hub gene rank | Hub gene symbol | Hub gene name | HIV modulation and overall impact on hub gene |
| --- | --- | --- | --- |
| 1 | ↑ EIF2AK2 * | Eukaryotic translation initiation factor 2 alpha kinase 2 | ▪ *Tat* shows multiple regulatory effects as: activates, inhibits, binds, is phosphorylated by, and is enhanced by EIF2AK2.<br>▪ Interaction with *Tat*, *Nef*, *gp120*, *gp160*, *Vpu*, *Vif*, *Pr55(Gag)*, and *capsid* |
| 2 | ↑ STAT2 * | Signal transducer and activator of transcription 2 | ▪ *Nef* induces phosphorylation, and *gp120* activates STAT2 |
| 3 | ↑↓ MAVS * | Mitochondrial antiviral signaling protein | ▪ *Nef* and *Vpu* block and degrade MAVS |
| 4 | ↓IFNAR1 * | Interferon alpha and beta receptor subunit 1 | ▪ *gp120* upregulates IFNAR1 |
| 5 | ↓IFNAR2 * | Interferon alpha and beta receptor subunit 2 | ▪ *Tat* upregulates IFNAR2 |
| 6 | ↑TRAF3 * | TNF receptor associated factor 3 | ▪ *Tat* downregulates TRAF3 |
| 7 | ↑SOCS3 | Suppressor of cytokine signaling 3 | ▪ *gp120* recruits SOCS3 while *Tat* both upregulates and downregulates SOCS3 |
| 8 | ↑BCL2L1 § | BCL2 like 1 | ▪ *Nef*, *gp120*, *Vpu*, *Vpr*, and nucleocapsid downregulate BCL2L1<br>▪ *Nef*, *Tat*, and *Vpr* also upregulate or interact with BCL2L1 |
| 9 | ↑ STING1 * | Stimulator of interferon response cGAMP interactor 1 | ▪ *Vpr* antagonizes STING |
| 10 | ↑CYCS * | Cytochrome c, somatic | ▪ *Tat*, *Vpr*, *gp120*, *gp160*, *retropepsin*, and *Nef* promote cytochrome c release<br>▪ *Tat*, *retropepsin*, and *Nef* also interact with or upregulate CYCS.<br>▪ *HIV1 replication* is enhanced by CYCS expression |

**Table 7.** Interactions Between Glioma Hub Genes and HIV Proteins.

| Hub gene rank | Hub gene symbol | Hub gene name | HIV modulation and overall impact on hub gene |
| --- | --- | --- | --- |
| 2 | ↑PIK3R1 *§ | phosphoinositide 3kinase regulatory subunit 1 | ▪ *Nef* activates the PI3K/Akt/mTOR pathway<br>▪ *gp120* activates PI3K and calcium mobilization<br>▪ *Tat* activates PI3K/Akt signaling<br>▪ *Vpr* stimulates PI3K/Akt/NFκB signaling in astrocytes |
| 3 | ↑↓IGF1R * | insulin like growth factor 1 receptor | ▪ *HIV1 gp120* upregulates IGF1 receptor (IGF1R) expression, along with multiple other pro-survival and apoptotic pathway genes, in human neuronal cells |
| 4 | ↑GRB2 * | growth factor receptor bound protein 2 | ▪ *Tat* interacts with GRB2.<br>▪ *Tat* promotes Ras activation via recruitment of GRB2, and **upregulates GRB2 isoform Grb33**, enhancing HIV1 LTR promoter activity.<br>▪ *HIV1 Pol and reverse transcriptase (RT)* physically interact with GRB2 in human cell lines |
| 5 | ↑↓PDGFB * | platelet derived growth factor subunit B | ▪ PDGFB inhibits *gp120* and *Tat*-induced neurotoxicity<br>▪ *gp120* and *Tat* upregulate PDGFB expression in endothelial cells |
| 6 | ↑TGFA * | transforming growth factor alpha | ▪ *Tat* upregulates transcription of the TGFA in an epidermal growth factor-dependent manner |
| 7 | ↑↓NRAS *§ | NRAS protooncogene, GTPase | ▪ *Nef* and *Tat* activate NRAS signaling linked to cellular compartments and kinase cascades<br>▪ *gp160* can both activate or inhibit<br>▪ NRAS supports **HIV replication** in T cells |
| 8 | ↑IGF1 * | insulin like growth factor 1 | ▪ IGF1 inhibits *gp120* by competing with its binding to the CD4 receptor<br>▪ IGF1 inhibits *Tat* mediated transactivation of the LTR promoter,<br>▪ *Tat* upregulates IGF1 and other antiapoptotic genes in stressed Kaposi's sarcoma cells.<br>▪ |
| 9 | ↑PIK3CB * | phosphatidylinositol 4,5bisphosphate 3kinase catalytic subunit beta | ▪ *Tat* and *Vpr* proteins manipulate the PI3K/Akt pathway to promote cell survival<br>▪ *gp120/gp160* engages chemokine receptors (CCR5/CXCR4)<br>▪ *Nef* activates the PI3K/Akt/mTOR pathway |

Hub gene regulation predicted as upregulation↑ or bidirectional ↑↓ in accordance with DE miRNA in pregnancy, \*
Genes annotated under cell protein kinase (Figure 7A). § *Shared with HCC. For the rank of hub genes in HCC,*
*refer to Figure 8C.*

## 4 Discussion

This study is unique in the longitudinal follow up of women before and after pregnancy events which is a strength of the study. Most longitudinal studies on pregnancy and peripheral blood miRNAs are related to birth outcomes and other pregnancies complications (20–22). By following women across these critical time points, our study provides a valuable opportunity to investigate the molecular shifts that occur during pregnancy and their potential implications for HIV susceptibility.

One of the several physiological adaptations during pregnancy involves the suppression of immune functions, irrespective of the women’s HIV status. These changes include early pregnant events such as reduction of complement events and cell-mediated immunity (23). In this sense, scientists and clinicians have yet to reach a consensus on whether pregnancy itself constitutes a risk factor to HIV acquisition (24–27). However, a recent study analyzing data from 2 large clinical trials concluded that the HIV risk increases during pregnancy and continuous through the first six months postpartum, with the highest risk occurring in the later stages of pregnancy (4). While association studies have laid the groundwork in exploring the link between pregnancy and HIV risk, too little is known, and limited empirical molecular evidence has been proposed.

MiRNAs are well-known immune modulators that regulate gene expression and, consequently, contribute to immune-related processes (28). It has been proposed that, in pregnancy, miRNAs are expressed in a coordinated manner from specific chromosomal clusters rather than individually (29, 30). Here, we determined changes in the serum miRNA across pre-pregnancy, pregnancy and breastfeeding reproductive states, providing evidence that molecular mechanisms governed by systemic miRNAs are also under physiological adaptation during those transitions. We tested the hypothesis that those individual miRNAs regulate pivotal genes and pathways related to the HIV interaction, influencing susceptibility during pregnancy.

Using a module-centric approach, MEA identified overrepresented functional groups and pathways, offering insights into coordinated biological processes and reducing the noise associated with individual gene analysis. In this context, modules were defined as clusters of functionally related genes based on shared gene ontology terms or pathway annotations, as computed by ClueGO’s kappa score-based grouping. Here we reported the enrichment of six pathways in pregnancy. Among those, the Hippo Signaling pathway stands out as physiologically relevant during pregnancy (31, 32), supporting the validity and biological relevance of our analysis. The Hippo Signaling pathway is functionally linked to the Cellular Senescence pathway (33), suggesting coordinated regulation. Cellular Senescence also shows physiological connections to the cancer-associated pathways— HCC and glioma — since altered cell cycle regulation is a hallmark of tumorigenesis (34, 35). The additional two pathways reflect overrepresentation of the adaptive immune response and viral infection processes, particularly those related to HSV-1. Curiously, a parallel has been proposed between maternal immune tolerance to the fetus during pregnancy and immune evasion mechanisms in cancer (36, 37). The idea is that the same immune system adaptations that help a fetus grow safely might also help cancer cells avoid being attacked. This might explain not only the cancer-related pathways found in this study, but also their linked genes. For instance, PDL-1 – a known cancer suppressor, predicted to be downregulated in our dataset – is one of the top hub genes found in Adaptive Immune Response term. Blood levels of PDL-1 were similar in pregnant women without cancer and in non-pregnant cancer patients, when compared to healthy non-pregnant women (37). Several genes contributing to the enriched pathways overlap with the HIV interactome. Therefore, our data highlights a convergence of molecular regulators that are targeted by miRNA DE in pregnancy that also share similarities with cancer pathways and may potentially be hijacked by HIV.

We identified a collection of 47 hub genes which are key proteins that play central roles within the networks of pregnancy-regulated set of miRNA targets. The miRNA-hub gene networks revealed a ‘one-to-many’ and ‘many-to-one’ relationships, shedding light on a complex regulatory landscape, in which individual miRNAs can influence multiple hub genes, while key hub genes may be co-regulated by several miRNAs. This reflects a coordinated regulatory control over critical biological processes. Moreover, the interaction of HIV proteins with hub genes suggests a potential viral strategy to hijack these key pathways, further disrupting host regulatory networks.

The global miRNA transcriptome during pregnancy identified also specific interactions between the hub gene targets, host proteins and HIV proteins in the HIV interactome. Among the host proteins, HLA-A showed the highest number of interactions, while Tat emerged as the central HIV modulator, alongside gp120 and Nef. HLA-A, predicted as upregulated, belongs to a gene complex that accounts for much of the observed variation in immune responses. It is highly polymorphic, and genetic variations have been demonstrated to be relevant in vertical transmission of HIV (38, 39). A large study of ∼9,800 HIV-infected individuals found that elevated HLA-A expression was associated with significantly higher viral load and lower CD4⁺ T-cell counts, indicating poorer virologic control (40). HIV Nef exhibits strong interaction with HLA-A and is known to downregulate its expression (41).

The co-targeting of hub genes by pregnancy-driven miRNAs and HIV proteins suggests convergent modulation, potentially representing a molecular *tug-of-war* between host regulation and viral interference throughout the course of pregnancy. This overlap may reveal potential points of susceptibility in pregnant women, where physiological adaptations to pregnancy intersects with viral strategies to hijack host pathways. Future studies should validate key pregnancy-associated miRNA interactions with HIV in relevant models and explore their timing throughout gestation. Integrating additional non-coding RNAs and transcriptomic data may uncover broader regulatory networks, open new possibilities in informed targeted HIV prevention strategies tailored to reproductive stages.

## 5 Conclusion

Global miRNA transcriptome analysis of longitudinal serum samples of women followed from preconception to pregnancy and postpartum breastfeeding identified pregnancy-regulated targets that work as hub genes per their intricate and high connection to other elements within identified protein networks. Some of these targets have known interactions with HIV modulators in a variety of models. Together, pregnancy-driven miRNAs and their targets may have a role in vulnerability and resistance to HIV and other viral infections.

## 6 Acknowledgment

Research reported in this publication was supported by the Eunice Kennedy Shriver National Institute of Child Health & Human Development of the National Institutes of Health under Award Number R01HD099091. The content is solely the responsibility of the authors and does not necessarily represent the official views of the National Institutes of Health.

## Notes

**Financial support and sponsorship:** Research reported in this publication was supported by the Eunice Kennedy Shriver National Institute of Child Health & Human Development of the National Institutes of Health under Award Number R01HD099091. The content is solely the responsibility of the authors and does not necessarily represent the official views of the National Institutes of Health.

### Competing Interest Statement

The authors have declared no competing interest.

